# Affinity Fine-Tuning of Boltz-2: An Open Framework for Protein-Ligand Potency Prediction in Drug Discovery

**DOI:** 10.64898/2026.05.26.727958

**Authors:** Samad Amini, Simone Sciabola, Ye Wang

## Abstract

Boltz-2 has enabled accurate binding affinity prediction by leveraging co-folded protein–ligand structures, but the absence of a public training recipe has limited its use in active drug discovery projects, where new experimental measurements and congeneric ligand series continually arrive during lead optimization. We present an open framework for affinity fine-tuning of Boltz-2, showing that adapting only the affinity prediction components with project-specific experimental data can make the model substantially more useful for lead optimization. We evaluate the approach in two internal studies: a multi-target retrospective benchmark against physics-based and machine learning baselines, and a large single-target study with up to 1,700 ligands. In both settings, fine-tuned Boltz-2 improves correlation over the off-the-shelf model, and in some cases reaches performance competitive with free energy perturbation (FEP) methods. By releasing this framework, we aim to enable the community to adapt Boltz-2 to the specific targets and data of their own drug discovery campaigns. The code is available at: https://github.com/molecularinformatics/Boltz2_affinity

## Introduction

Accurate prediction of small-molecule binding affinity remains a central problem in hit identification and, especially, lead optimization. For more than a decade, relative binding free energy methods such as FEP+ have served as the gold standard (Wang et al., 2015; Abel et al., 2017). However, their reliance on molecular dynamics and carefully prepared perturbation maps makes them computationally expensive and ill-suited to the throughput demands of modern discovery campaigns. The emergence of co-folding models — beginning with AlphaFold3 (Abramson et al., 2024) and continued in the open-source Boltz series (Wohlwend et al., 2024; Passaro et al., 2025) — has reshaped this landscape by jointly predicting protein–ligand complex structure at a fraction of the cost. Boltz-2 extends this line of work with a dedicated affinity head and has approached FEP+ accuracy on the public FEP+ benchmark while being orders of magnitude faster.

Unlike physics-based free energy methods, which depend on fixed functional forms and expensive conformational sampling, co-folding models learn protein–ligand representations directly from data and can therefore be adapted as new experimental measurements are collected. This is particularly valuable in lead optimization, where potency data accumulates over congeneric or closely related series and encodes local SAR that may be poorly captured by a general-purpose model. Although Boltz-2 learns broad protein–ligand patterns from public data, its off-the-shelf form cannot leverage the project-specific information contained in internal assay measurements. Fine-tuning addresses this limitation by adapting the affinity head to the target, chemotype, and assay context of an active program, improving specialization to the chemical space that matters most for that project.

In this work, we present an open framework for affinity fine-tuning of Boltz-2 that makes project-specific adaptation practical and reproducible. Our implementation extends the public model with the components needed for supervised affinity tuning and is designed to be accessible for researchers who want to adapt structure-based foundation models to their own datasets. We then evaluate the framework on two internal drug discovery studies. In the first, we extend a previously published retrospective benchmark across four targets and five congeneric series, positioning fine-tuned Boltz-2 alongside physics-based free energy methods and other ML baselines. In the second, we fine-tune on up to 1,500 ligands from a single internal program and measure the resulting improvement in correlation against the public checkpoint. Together, these studies demonstrate that lightweight fine-tuning of the affinity head on project-scale potency data is sufficient to move Boltz-2 from a strong general-purpose predictor to a tool competitive with FEP on the targets that matter to a given campaign.

## 2. Related Works

Physics-based methods such as FEP+ (Wang et al., 2015) remain an important standard for ranking congeneric ligands in lead optimization, but their molecular-dynamics-based sampling and setup costs limit routine use at large scale. By contrast, MM/GBSA (Genheden & Ryde, 2015) is substantially cheaper and widely used, although its accuracy is more system dependent. Early structure-based deep learning methods such as KDeep (Jiménez et al., 2018) showed that 3D convolutional networks can learn affinity directly from protein–ligand complex representations, and DeltaDelta neural networks (Jiménez-Luna et al., 2019) further emphasized lead-optimization settings by learning pairwise potency differences within congeneric series. More recent work has shifted toward graph-based and geometry-aware architectures, including interaction-focused GNNs such as PIGNet (Moon et al., 2022) and equivariant representations like Uni-Mol (Zhou et al., 2023), as well as diffusion-based models such as DiffDock (Corso et al., 2023) that learn protein–ligand pose distributions and expose confidence signals that can be repurposed for ranking. In parallel, co-folding models have begun to unify structure and affinity prediction in a single framework: AlphaFold3 (Abramson et al., 2024) established high-accuracy joint protein–ligand structure prediction without a dedicated affinity output, and Boltz-2 (Passaro et al., 2025) extends his paradigm to joint structure and affinity prediction. Our work builds on this trajectory by focusing on fine-tuning Boltz-2 for project-specific affinity prediction in realistic medicinal chemistry settings.

## 3. Methods

We extend the public Boltz-2 codebase with four components to enable efficient project-specific affinity fine-tuning: a regression loss, a parameter freezing strategy, a precomputed embedding pipeline, and affinity-aware data processing. The full implementation is released as an open framework built on top of the Boltz-2 training infrastructure. Rather than re-training the full co-folding model, our framework is designed around lightweight adaptation of the affinity prediction components while keeping the rest of the network fixed.

Boltz-2’s affinity module produces continuous predictions (pIC50) via a regression head. We train the affinity head with a Huber loss (*δ* = 0.5), which behaves quadratically for small residuals and linearly for large ones. This makes it more robust than mean-squared error to the heavy-tailed residuals typical of internal potency assays, where a handful of mis-measured or outlier compounds can otherwise dominate the gradient. Given a predicted affinity *ŷ* and experimental value *y* on the *log*10(IC50), derived from an IC50 measured in *µM*, the loss is

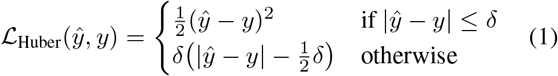

with transition point *δ* treated as a fixed hyperparameter. We also implemented pairwise ranking and focal binary classification losses, but set their weights to zero in all reported experiments, training exclusively on continuous affinity regression.

Efficient fine-tuning is achieved by freezing all Boltz-2 parameters except the affinity prediction head. An `affinity only finetuning` flag restricts gradient updates to three affinity modules (`affinity module`, `affinity module1`, `affinity module2`), reducing trainable parameters from 515M (full model) to 8.7M. All frozen modules are set to `.eval()` mode to disable stochastic operations including dropout. This enables fine-tuning on a single GPU within hours.

A central practical challenge in affinity fine-tuning is that repeatedly recomputing frozen trunk representations is unnecessarily expensive when only the affinity head is being updated. To address this, we implemented a two-stage workflow. First, the full Boltz-2 model is run once over the dataset to precompute and store the single and pair representations for each example. Second, during fine-tuning, these cached representations are loaded directly and used in place of a fresh forward pass through the frozen trunk and structure modules. Because affinity fine-tuning uses ligand-centered cropping, the cached pair representation must be remapped to the cropped token subset; we handle this by tracking the original token indices during preprocessing and re-indexing the precomputed pair tensor accordingly at fine-tuning time. This design substantially reduces the computational cost of fine-tuning and makes project-level adaptation practical on modest hardware.

Each training record is annotated with an `AffinityInfo` object storing the continuous affinity value, a binary binder label (unused here), an assay identifier (for future pairwise training), and molecular weight. Annotations are parsed from YAML input files and propagated through the featurizer to produce per-sample ground-truth labels.

## 4. Experiments

### 4.1. Study 1: Multi-Target Retrospective Benchmark

We benchmark on the dataset introduced by Bansal et al. (2024), comprising 172 compounds across four internal protein targets and five congeneric series. This benchmark includes one enzyme target and three kinase targets, with two separate datasets for one kinase target reflecting different medicinal chemistry optimization regimes. We use the same train/test splits and Tanimoto similarity profiles reported in the original study (Table 1 of Bansal et al. (2024)), with per-target training sets ranging from 52 to 195 compounds. This setting is useful because it spans chemically realistic project series and includes cases that are known to be more challenging for structure-based methods.

We extend this benchmark by adding two Boltz-2 conditions to the original comparison: Boltz-2 (Default), corresponding to the public checkpoint used without project-specific adaptation, and Boltz-2 (Fine-tuned), obtained by fine-tuning the affinity components on project data using the framework described above. We compare these against a compact set of representative baselines: FEP+ as the strongest physics-based reference in the original study, DeltaDeltaG, and KDEEP in both default and retrained forms. Following the prior benchmark, performance is measured using the Pearson correlation coefficient between predicted and experimental binding affinities, reported separately for each of the five datasets and summarized in Figure 1. The original benchmark showed that FEP+ was the strongest physics-based baseline overall, while retrained machine-learning models substantially outperformed their off-the-shelf versions, motivating this same adaptation-based evaluation for Boltz-2.

**Figure 1.**
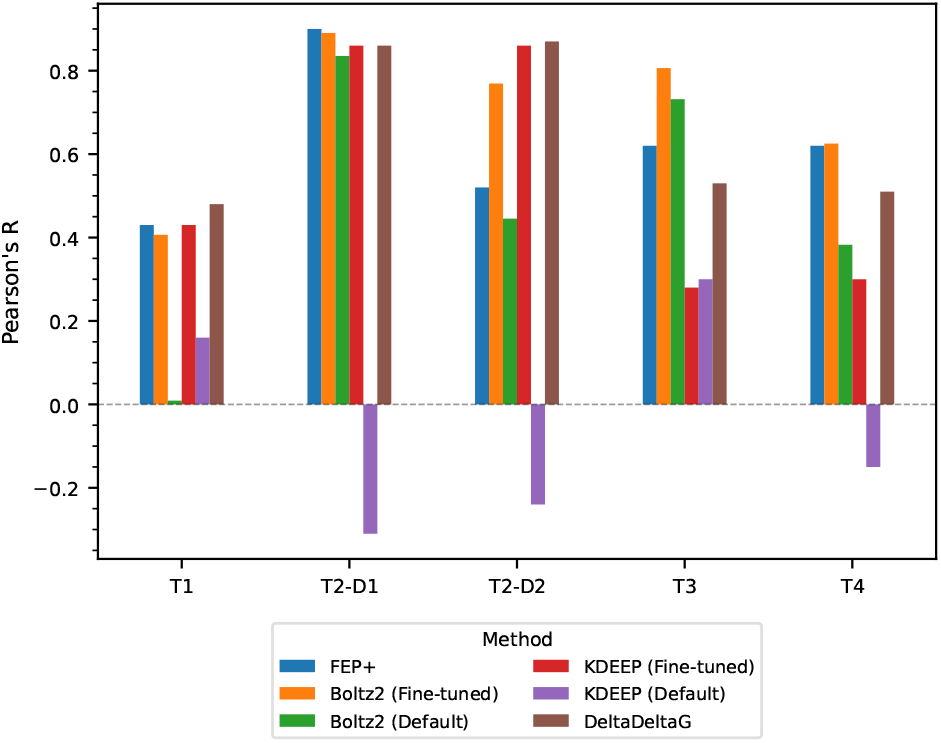
Pearson correlation obtained by correlating experimental binding affinity with the predicted potencies for each target across various methods shown in the legend. T1–T4 denote Target 1–Target 4, with T2 split into two independent datasets (T2-D1 and T2-D2)

### 4.2. Study 2: Single-Target Time-Split Evaluation

We evaluate on a single internal program with longitudinal experimental data, where compounds were synthesized and assayed in monthly batches over approximately two years.

This setting mirrors real-world lead optimization, where a model must generalize to structurally novel compounds arriving after the training cutoff. We define 19 time splits (*t*_5_– *t*_23_), where each split *t*_*k*_ uses all compounds from batches 1 through *k−* 1 as training data and batch *k* as the held-out test set. Training set size grows from roughly 276 to 1,714 compounds across the splits, reaching up to 1,734 total train/test ligands at the last time split.

At each split, Boltz-2 (Fine-tuned) is retrained from the public checkpoint using only the data available up to *t*_*k−*1_. We compare against Boltz-2 (Default), KDEEP (retrained at each split on the same training data), and Docking (Glide SP, static baseline with no retraining). All methods are evaluated by Pearson’s *R* on the held-out batch. Figure 2 shows Pearson’s *R* for all four methods across the 19 time splits. Boltz-2 (Default) provides a weak but nonzero baseline throughout (mean *R* = 0.38), confirming that the public checkpoint transfers some signal to this target without any adaptation. Fine-tuning substantially and consistently improves over the default checkpoint across nearly all splits (mean *R* = 0.76), with particularly large gains in the mid-to-late splits as training data accumulates. Boltz-2 (Fine-tuned) matches or exceeds docking on the majority of splits and outperforms KDEEP on all but two. The improvement over the default checkpoint is relatively stable even at early splits, suggesting that the affinity head adapts efficiently to limited data when the trunk representation is fixed.

**Figure 2.**
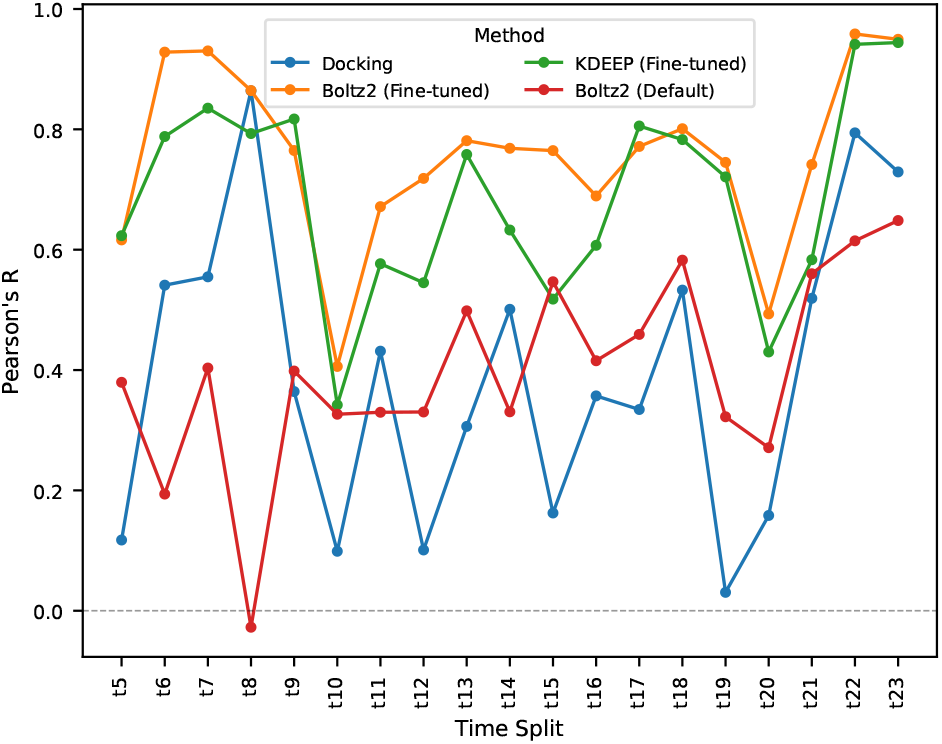
Pearson’s *R* across 19 monthly time splits (*t*_5_–*t*_23_) on a single internal target. At each split, models are retrained on all prior batches and evaluated on the held-out batch.

## 5. Analysis

Across both studies, the gap between Boltz-2 (Default) and Boltz-2 (Fine-tuned) is large and consistent. In Study 1, the default checkpoint provides a modest but nonzero signal on most targets (Figure 1), confirming that the public model transfers useful structural representations to unseen targets without any adaptation. However, fine-tuning on as few as 50–200 project-specific compounds produces gains of 0.2– 0.4 Pearson’s *R* units across targets, bringing the model into the range of FEP+ and retrained ML baselines. In Study 2, this pattern holds longitudinally: Boltz-2 (Default) hovers around *R≈* 0.38 throughout the timeline, while Boltz-2 (Fine-tuned) averages *R≈* 0.74 across 19 splits (Figure 2), demonstrating that adaptation from the public checkpoint is effective even in the early splits where training data is limited.

On the multi-target benchmark, Boltz-2 (Fine-tuned) matches or exceeds FEP+ on four of five datasets (T2-D1, T2-D2, T3, T4), and on T2-D2 specifically outperforms FEP+ despite that dataset being characterized by P-loop flexibility that is known to challenge physics-based sampling methods (Bansal et al., 2024). This suggests that structure-based fine-tuning may implicitly capture some of the conformational dependence that FEP+ addresses through explicit simulation. On T1 — an enzyme with a large hydrophobic pocket and multiple binding modes — Boltz-2 (Default) provides near-zero correlation (*R≈* 0.01), but fine-tuning recovers substantial signal (*R≈* 0.40), approaching FEP+ (*R ≈* 0.43). The remaining gap relative to DeltaDeltaG on this target may reflect the difficulty of the binding site rather than a limitation of the fine-tuning approach, as T1 was the most challenging target across all methods in the original benchmark (Bansal et al., 2024). In Study 2, fine-tuned Boltz-2 tracks closely with Fine-tuned KDEEP across the full timeline and pulls ahead on several later splits as training data accumulates.

KDEEP (Default) shows strongly negative Pearson’s *R* on three of five benchmark targets, and Boltz-2 (Default) provides only weak signal in Study 2 despite access to co-crystal structures. This confirms a well-established observation in the field (Bansal et al., 2024): ML methods trained on public data generalize poorly to project-specific assay conditions, chemical series, and target biology. The practical implication is that off-the-shelf use of any ML affinity model including structure-based models like Boltz-2 is unlikely to be reliable for active lead optimization campaigns without project-specific adaptation.

### Impact Statement

This paper advances the field of Machine Learning for drug disovery by providing an open-source framework for fine-tuning the Boltz-2 affinity module. By releasing the implementation where the or
iginal training code remains unavailable, our work enables broader community access to high-fidelity molecular modeling tools, potentially accelerating the identification of therapeutic candidates.

## Notes

### Competing Interest Statement

The authors have declared no competing interest.

